# Phytochromes mediate germination inhibition under red, far-red, and white light in *Aethionema arabicum*

**DOI:** 10.1101/2022.06.24.497527

**Authors:** Zsuzsanna Mérai, Fei Xu, Andreas Musilek, Florian Ackerl, Sarhan Khalil, Luz Mayela Soto-Jiménez, Katarina Lalatović, Cornelia Klose, Danuše Tarkowská, Veronika Turečková, Miroslav Strnad, Ortrun Mittelsten Scheid

## Abstract

The view on the role of light during seed germination stems mainly from studies with *Arabidopsis*, where light is required to initiate this process. In contrast, white light is a strong inhibitor of germination in other plants, exemplified by accessions of *Aethionema arabicum*, another *Brassicaceae.* Their seeds respond to light with gene expression changes of key regulators converse to *Arabidopsis*, resulting in antipodal hormone regulation and prevention of germination. The photoreceptors involved in this process in *A. arabicum* were unknown. Screening the first mutant collection of *A. arabicum*, we identified *koy-1*, a mutant that lost light inhibition of germination, due to a deletion in the promoter of *HEME OXYGENASE 1*, the gene for a key enzyme in the biosynthesis of the phytochrome chromophore. *koy-1* seeds are unresponsive to red- and far-red light and hyposensitive under white light. Comparison of hormone and gene expression between wild type and *koy- 1* revealed that very low light fluence stimulates germination, while high irradiance of red and far-red light is inhibitory, indicating a dual role of phytochromes in light-regulated seed germination. The mutation also affects the ratio between the two fruit morphs of *A. arabicum*, suggesting that light reception via phytochromes can fine-tune several parameters of propagation in adaptation to conditions in the habitat.

**One sentence summary:** Characterization of a phytochrome chromophore biosynthesis mutant demonstrates an active role of phytochromes in the light-inhibited seed germination in *Aethionema arabicum*.

## Introduction

Seed germination, a bottleneck in the plant life cycle, is controlled by environmental factors like temperature, humidity, nitrate, and light (reviewed in Vleeshouwers et al., 1995; Finch-Savage and Leubner-Metzger, 2006). Light quality and quantity provide information to the seed about the coverage by soil, the canopy shade, or day-length (Pons, 2000; Chen et al., 2014). Due to experiments with the classical model plants in germination research, like Arabidopsis or lettuce, light has long been considered to alleviate dormancy and stimulate germination, and the molecular basis of this regulation has been extensively studied (Vleeshouwers et al., 1995). The requirement for light induction to seed germination has been considered as a dept-sensing mechanism of small-seeded plants to avoid germination deep underground if the resources would not allow them reaching the surface (Seo et al., 2009). However, seed germination response to white light can also be neutral, or even negative, when white light inhibits the process (Takaki, 2001; Yang et al., 2020). Photoinhibition of germination, termed negative photoblasty, has been observed in several, phylogenetically distant taxa across angiosperms (Koller, 1957; Botha and Small, 1988; Thanos et al., 1991; Thanos et al., 1994; Carta et al., 2017; Vandelook et al., 2018; Mérai et al., 2019). White light was found to cause delay or partial inhibition of germination in tomato variants, while it strongly inhibits the germination of *C. lanatus* seeds (Yaniv and Mancinelli, 1968; Botha and Small, 1988). The light-inhibited germination is interpreted as an adaptive trait to avoid germination on the surface in open and arid habitats like coastal dunes, deserts, shrublands, or grasslands (Pons, 2000; Carta et al., 2017). *Aethionema arabicum*, belonging to the *Brassicaceae* and originating from semi-arid open habitats, provides an opportunity to study the largely unknown molecular basis of negative photoblasty. Some accessions of this species have light-neutral seed germination, but seed germination of an accession from Cyprus (CYP) is strongly inhibited by continuous white light and long-day conditions, while seeds germinate under short days despite strong light illumination (Mérai et al., 2019). We speculated that the light-inhibited germination might be a day-length measuring mechanism to ensure the proper timing of germination at early spring, matching the germination temperature optimum at 14°C, and proposed *A. arabicum* as a suitable model plant to study the rewiring of the light response in seeds (Mérai et al., 2019). The germination of CYP seeds is strongly inhibited by white, red, far-red, and blue light, indicating the involvement of phytochromes and possibly other photoreceptors in the process (Mérai et al., 2019). However, the specific photoreceptors mediating germination inhibition in *A. arabicum* are unknown. Assuming that loss of genes encoding signaling or other regulatory components would result in an easily scoring phenotype, we generated the first mutant collection in *A. arabicum* and screened for germination of light-exposed seeds. Not surprisingly, one of the first mutations identified confirmed an important role for a known component in light reception, but it added an unexpected facet, explained the negative photoblasty in the wild type, and revealed novel links to developmental parameters.

Plants perceive and transduce light signals through photoreceptors. Red and far-red spectra (600-750 nm) are received by phytochromes (reviewed in Gyula et al., 2003; Li et al., 2011). UV-A and blue light (320-500 nm) perception is mediated by three flavin-based photoreceptor families, the cryptochromes (crys), phototropins (phots) and the Zeitlupe family (ztl, fkf1 and lkp2) (reviewed in Christie et al., 2015). Additionally, UV-B light (282- 320 nm) is perceived by UV RESISTANCE LOCUS 8 (UVR8) (Rizzini et al., 2011). Blue light can be involved in the control of dormancy induction in cereals via cryptochrome CRY1 and in the control of dormancy alleviation in *Arabidopsis* via phyB (Shropshire et al., 1961; Barrero et al., 2014; Stawska and Oracz, 2019). However, the connection between light and seed germination is most prominent for longer wavelengths perceived by phytochromes, extensively studied for decades (Casal and Sanchez, 1998; Yang et al., 2020).

The light-sensing chromophore of all phytochromes in higher plants is an open-chain tetrapyrrole called phytochromobilin (PφB). It is synthetized in the plastids from 5- aminolevulinic acid in a series of enzymatic reactions including an oxidative cleavage of a heme intermediate into biliverdin IX by a ferredoxin-dependent heme oxygenase (HO). Biliverdin is reduced to PφB by the PφB synthase. The chromophore is then exported to the cytosol and assembles autocatalytically with the phytochrome proteins, forming the photoreversible holophytochromes (Brown et al., 1990; Mahawar and Shekhawat, 2018). Arabidopsis has five phytochromes that play a role in diverse signaling pathways, e.g., regulating seed germination, seedling de-etiolation, shade avoidance, flowering time and hormonal metabolism. In darkness, phytochromes are synthetized in the inactive P_r_ form that converts to the active P_fr_ form upon absorbing red light (660 nm). The active P_fr_ form initiates the signaling cascade leading to the phytochrome response (Furuya and Schäfer, 1996). P_fr_ is inactivated quickly upon far-red light (730 nm) absorption or by spontaneous relaxation via the slower dark reversion (Mancinelli, 1994; Quail, 1997). The light-labile phyA is the only receptor that perceives and mediates responses in far-red light, while phyB-phyE are considered light-stable red-light receptors, with a predominant role of phyB (Dehesh et al., 1993; Nagatani et al., 1993). Much insight into their function stems from phytochrome-deficient mutants, in which either the apoproteins or the chromophore are missing. Single *phya*, *phyb*, etc. mutants allow the functional characterization of the individual phytochromes (Koornneef et al., 1980; Quail et al., 1995), whereas mutations affecting the chromophore biosynthesis pathway result in a loss or severe reduction of all photoreversible phytochromes and are impaired in photomorphogenesis (reviewed in Terry, 1997). This was shown for mutants in several plant species lacking heme oxygenase and PφB synthase (Koornneef et al., 1980; Koornneef et al., 1985; Chory et al., 1989; Kraepiel et al., 1994; Lamparter et al., 1996; van Tuinen et al., 1996; Weller et al., 1996; Weller et al., 1997; Izawa et al., 2000; Sawers et al., 2004). Here we present *koy-1*, a chromophore-deficient heme oxygenase mutant in *Aethionema arabicum* that has lost the inhibition of seed germination under red, far-red, and white light and shows a shifted fruit morph ratio, revealing novel phenotypes never observed in chromophore mutants of other species.

## Materials and Methods

### Plant material

Experiments were conducted with *Aethionema arabicum* (L.) Andrz. ex DC. accessions TUR ES1020 and CYP (obtained from Eric Schranz, Wageningen). WT and *koy-1* plants were propagated for seed material under 16 h light/19°C and 8 h dark/16°C diurnal cycles, under ∼300 μmol m^-2^ s^-1^ light intensity. Fruits and seeds were counted manually after complete maturation from 6 or 8 plants, under 200 or 300 μmol m^-2^ s^-1^ light intensity, respectively. WT and *koy-1* plants were randomly distributed on the shelves.

### Plant chambers and light source

The mutant screen and white light germination assays were carried out in a Percival plant growth chamber equipped with fluorescent white light tubes with a spectrum from 400- 720 nm. Red, far-red, and blue light treatments were performed using LED light sources, peaking at 658 nm, 735 nm and 450 nm, respectively. For experiments under higher light intensities (Fig. 1A,F) a Percival growth chamber was equipped with a Valoya LightDNA-8 LED light source (https://www.valoya.com/lightdna/). Light spectra and intensity were measured by LED Meter MK350S (UPRtek).

**Figure 1.**
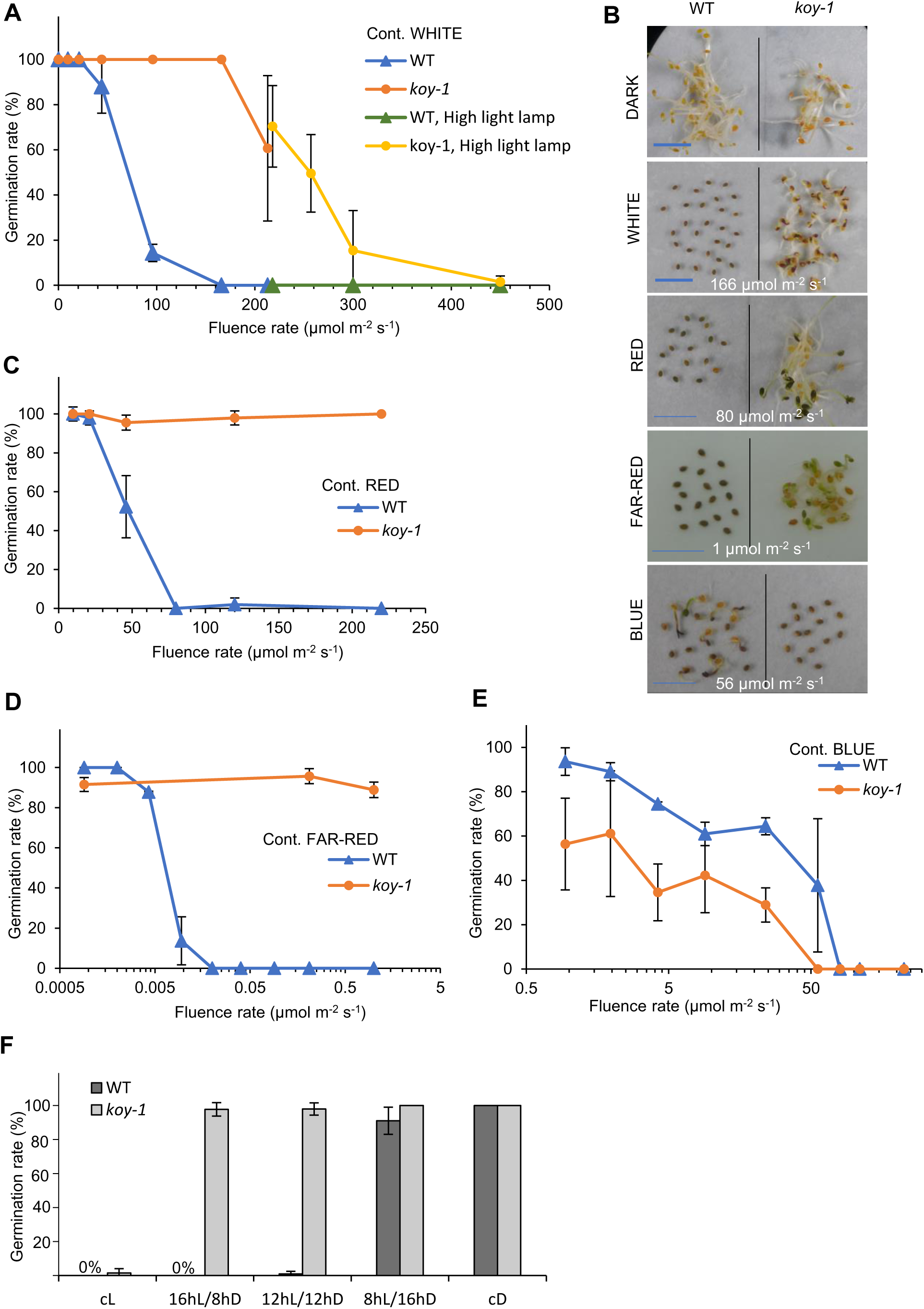
Germination response of WT and *koy-1* mutant to light intensity and light color. Germination of WT and koy-1 seeds after six days under different intensities of continuous white (A,B), red (B,C), far-red (B,D), and blue (B,E) light. (F) Diurnal regulation of seed germination under 450 µmol m-2 s-1 white light. Daily dark and light regime is indicated as hour D and L. cL and cD indicate constant light and dark, respectively. Error bars indicate standard deviations of three biological replicates.

### *Aethionema arabicum* genome and annotations

*Aethionema arabicum* genome version 3.0, gene models, cDNA, and protein annotations (version 3.1) were obtained from *A. arabicum* database (https://plantcode.cup.uni-freiburg.de/aetar_db/) (Haudry et al., 2013; Fernandez-Pozo et al., 2021). Accession numbers used in this study are listed in Supplementary Table S1.

### Fast neutron mutagenesis and screening

Seeds of *A. arabicum* (CYP accession) were irradiated in a TRIGA Mark-II research reactor in the Atomic Institute of the Technical University, Vienna (https://ati.tuwien.ac.at/startpage/EN/). Two thousand seeds were irradiated with either 1 kW or 5 kW for one minute, corresponding to 8x10^9^ cm^-2^s^-1^ and 4x10^10^ cm^-2^s^-1^ thermal neutron flux density, respectively. Seeds were plated for germination on the day of irradiation on wet filter paper and kept in darkness at 14°C for 6 days. Approximately 2000 individually bar-coded M1 seedlings were planted in a glasshouse under 19°C and 8 h dark/16°C diurnal cycles. One thousand three hundred twenty plants produced seeds representing the M2 mutant seed bank. Lines with seed amounts insufficient for direct screening were further propagated to M3 seed lots by pooling the seeds from five M2 plants. Seeds were subjected to the forward genetic screen not earlier than six months after harvest. Germination was scored after 7 days under continuous white light illumination with 160 µmol m^-2^ s^-1^ light intensity at 14°C in a growth chamber.

### Identification of the causative mutation in *koy-1*

*koy-1* originates from a single seedling that germinated under light from one of the M2 seed batches originating from the lower, 1 kW radiation. From the same plate, eight other seeds were germinated in darkness, propagated and grown in parallel with *koy-1*. A wild-type phenotype in all their progeny and the mutant phenotype of *koy-1* progeny were confirmed for seeds of two following generations. These seed pools originated from the same M1 mother plant and were expected to share many of the non-causative mutations, except the causative mutation of *koy-1*. Eighteen koy-1 seedlings versus 64 wild type seedlings were pooled for DNA isolation and sequenced on an Illumina H2500 platform with 100 bp single end mode by the Next Generation Sequencing Facility of the Vienna BioCenter Core Facilities (VBCF), member of the Vienna BioCenter (VBC), Austria. Sequencing reads were processed by the CLC Genomics Workbench 9.5.1 software (Qiagen). Reads were mapped to the *Aethionema arabicum* PacBio contigs originating from the CYP accession (see below). After removal of duplicated reads, 55.8 and 48.1 million reads resulted in 21.98x and 18.97x coverage in the *koy-1* and wild-type pool, respectively. We expected the *koy*-1 pool to be homozygous for the mutation and the wild-type pool homozygous for the reference genome sequence, therefore variants were called with 90% minimum frequency in the *koy*-1 pool and filtered against the wild-type reads. Seventeen single nucleotide polymorphisms were uniquely found in the *koy-1* pool, but none of them overlapped with genic regions. InDel variants were called if evidence for insertion or deletion was detected in a minimum of six reads and if they appeared as homozygous variants (variant ratio >0.8) in the *koy-1* pool but not in the wild-type pool. Out of the 63 variants that fulfilled these criteria, two overlapped with genic regions. One of them was sorted out due to mapping errors, while the remaining one corresponded to the promoter deletion of the HEME OXYGENASE 1 (*Aa31LG4G11055*). The deletion was confirmed by PCR with the primers 5′CCTGGTGGTGGTAATGAACTC and 5′CGGTGGGGCAAAAGCGAATCC. The promoter sequence was analyzed using the PlantCARE tool (Lescot et al., 2002).

### Germination test

After seed harvest, seed stocks were kept dry at 24°C for six months, except for the experiments in Fig. 7B,C where eight-week-old semi-dormant seed batches were used to test the germination induction of light pulses. One seed batch consists of the harvest from at least 6 plants; replicates represent different seed batches. Except that in Supplemental Fig. S1, all germination tests were conducted at the optimal temperature of 14°C for six days in Petri dishes on 2-layer filter paper wetted with distilled H_2_O and supplemented with 0.1% plant preservative PPM (Plant Cell Technology). For dark treatments, seeds were placed on wet filter paper under complete darkness. Segregation assays were performed under 120 µmol m^-2^ s^-1^ white light.

### Hypocotyl elongation test

Seeds were plated on Petri dishes with 2-layer filter paper wetted with distilled H_2_O and supplemented with 0.1% plant preservative PPM (Plant Cell Technology). To induce germination, plates were kept in dark at 14°C for two and a half days, then transferred to red, far-red, or blue light, or kept in darkness as a control. After five days, seedlings were transferred to 1% (v/w) agarose plates and images were captured. Images were analyzed using Fiji software (https://fiji.sc/).

### Chlorophyll and anthocyanin measurement

As described for the hypocotyl elongation test, seeds were induced for germination in darkness followed by a 5-day-long light treatment or kept in darkness. Chlorophyll and anthocyanin measurements were performed as described (Chory et al., 1989; Holm et al., 2002), using 100 mg seedlings harvested in darkness.

### Quantitative RT-PCR

WT and *koy-1* seeds were illuminated at 14°C with 80 µmol m^-2^ s^-1^ red light, 1 µmol m^-2^ s^-1^ far-red light, under the condition where the WT seed germination is fully inhibited. The intensity of blue illumination was chosen to be 56 µmol m^-2^ s^-1^, where the germination is fully inhibited for *koy-1* mutant but not for WT seeds. As a control, seeds were kept in darkness. After 24 h exposure, seeds with intact seed coats were collected for RNA extraction, with three biological replicates for each sample. RNA extraction, cDNA synthesis and quantitative qPCR were performed as described (Mérai et al., 2019), using the primer pairs listed in Supplementary Table S2. The geometric mean of Aethionema orthologues of POLYUBIQUITIN10 (*AearUBQ10, Aa3LG9G835*) and ANAPHASE-PROMOTING COMPLEX2 (*AearAPC2*, *Aa31LG10G13720*) was used for normalization (Mérai et al., 2019). For each gene, the expression levels are presented as fold change relative to the level of the dark samples in WT seeds, where the average expression was set to one. Statistical analysis was done using the SATQPCR tool (Rancurel et al., 2019). Error bars represent standard deviation. Asterisks indicate significant differences from the WT or *koy-1* dark level with p- values as *p<0.05, **p<0.01, ***p<0.001, and ****p<0.0001 calculated with the Tukey test.

### Measurement of hormone levels

For ABA and GA analysis, seed samples were collected as described for RNA extraction, except that five biological replicates were prepared per sample. Sample preparation and targeted profiling of ABA (Turecková et al., 2009) and the members of GA biosynthetic as well as metabolic pathways (Urbanová et al., 2013) were performed as described earlier (Mérai et al., 2019).

### Extraction of high molecular weight DNA

The high molecular weight DNA was extracted as described in (Hofmeister et al., 2020; Barragan et al., 2021) with minor modifications. Twenty grams of leaf tissue were harvested from 4-week-old plants grown under standard conditions and ground in liquid nitrogen. The fine powder was resuspended in 200 ml cold nuclear isolation buffer (10 mM Tris-HCl, 100 mM KCl, 10 mM EDTA pH 8.0, 0.5 M sucrose, 4 mM spermidine, and 1 mM spermine) and filtered through two layers of miracloth. After adding Triton X-100 to 1% v/v final concentration, the nuclei were pelleted at 2,200 x *g* at 4°C for 15 min and resuspended in 40 ml nuclear isolation buffer with 1% v/v Triton X-100. Nuclei were lysed by addition of G2 lysis buffer (Qiagen) supplemented with 50 µg/ml RNaseA and 200 µg/ml Proteinase K. Samples were incubated first at 37°C for 30 min and then at 50°C for 2 h. Nuclear debris was pelleted by centrifugation at 10,000 x *g* at 4°C for 15 min. The supernatant was further purified using the Blood&Cell Culture DNA kit (Qiagen) according to the manufacturer′s protocol. After elution, the DNA was precipitated overnight at 4°C by adding 0.7 volume of isopropanol. The visible DNA pellet was spooled out using a glass rod, transferred to 300 µl 1xTE buffer and completely dissolved. The DNA size and integrity were assayed using the Femto Pulse System (Agilent) and confirmed as a single 165 kb-large intact band.

### PacBio sequencing

Library preparation and PacBio sequencing were performed by the Next Generation Sequencing Facility at Vienna BioCenter Core Facilities (VBCF), member of the Vienna BioCenter (VBC), Austria. The DNA was sheared to 34-165 kb and size-selected by a BluePippin instrument (sage science) with a lower cutoff at 25 kb. The library was prepared using the SMRTBell express Kit (PacBio) and sequenced on a PacBio Sequel platform (PacBio) with a yield of 22.04 Gb. This acquired 816,659 subreads with 14.7 kb mean length, 4.28 kb lowest quartile length, and 22.5 kb highest quartile length, resulting in 12,063,792,222 total base pairs.

### Assembly and annotation of PacBio contigs

The genome was assembled from 22 Gb of Sequel long read data using *Canu* (version 1.8; genomeSize = 235 m; corMaxEvidenceErate = 0.15, other parameters in default settings) (Koren et al., 2017). Raw contigs were polished with *Arrow* in two rounds using the default parameters and Pacbio reads. Mapping and alignment of cDNAs generated with GMAP (Wu and Watanabe, 2005), using the *A. arabicum* cDNA annotations version 3.1 (Fernandez-Pozo et al., 2021). Assembled PacBio contigs are deposited at X (during review).

### *In vivo* spectroscopy

The total amount of photoreversible phytochrome was measured by dual-wavelength ratiospectrophotometer (ratiospect) as described in (Klose, 2019), using seedlings of WT and *koy-1* that had been germinated on wet filter paper at 14°C in darkness for six days and collected under safety green light.

## Results

### Screen for irradiation-induced mutants with hyposensitive seed germination in response to white light

The strong inhibition of seed germination in an *Aethionema arabicum* accession originating from Cyprus (CYP) by light, independent of wavelength (Merai et al., 2019), allowed to characterize this feature with a straightforward genetic approach. Defects in light perception or signal pathways to the hormonal control of the inhibition should result in seeds able to germinate under light, and the respective mutations are expected to help identifying important components of the regulation. Therefore, we prepared the first mutant collection of *Aethionema arabicum* in the background of the CYP accession as wild type (WT). As the thick seed coat and mucilage would limit the uptake of chemical mutagens, we irradiated seeds with fast neutrons, planted them individually and collected their seeds, representing the M2 generation. At least 30 seeds per lines were assayed at optimal temperature (14°C) for germination under 160 µmol m^-2^ s^-1^ light intensity, conditions under which less than 1% of the WT seeds germinate. Among approximately 1300 lines screened, we found four well-established lines as primary mutant candidates. These were further propagated for two generations to confirm the heritability of the mutant phenotype. In the following, we present the first mutant, for which we identified the genetic defect and the possible role of the gene product. We named the mutant *koyash-1* (*koy-1*), after the god of sun in Turkic mythology.

While wild type seed germination is already strongly reduced at white light intensity of 100 µmol m^-2^ s^-1^, *koy-1* seeds germinate at a rate close to 100% at up to 166 µmol m^-2^ s^-1^ (Fig. 1A). Using a set-up extending the intensity range, *koy-1* seeds were fully inhibited only at 450 µmol m^-2^ s^-1^ (Fig. 1A). Wild-type and *koy-1* germinate over 90% in darkness (Fig. 1A,B), excluding differences in germination potential between the seed batches. The optimal temperature for seed germination with an optimum range between 11-14°C and low germination rate at 8°C or 20°C, is also similar between WT and *koy-1* (Supplemental Fig. S1), with a slightly lower germination rate at 17°C. Therefore, the mutation in *koy-1* changes only the response to light, not to seed vigor or temperature.

### Light spectra specificity of the *koy-1* mutant phenotype

The germination of wild-type seeds is strongly inhibited by continuous red, far-red, and blue light in a dosage-dependent manner (Fig. 1B-E). The effective intensity of blue and red light is comparable: germination is strongly inhibited at 50 µmol m^-2^ s^-1^ and completely abolished at 80 µmol m^-2^ s^-1^ and above (Fig. 1C,E). Interestingly, the inhibition by far-red light is four orders of magnitude stronger, resulting in complete inhibition at and above 0.02 µmol m^-2^ s^-1^ intensity (Fig. 1D). *koy-1* seeds could completely germinate under red and far-red illumination even up to very high intensities (220 or 1 µmol m^-2^ s^-1^, respectively). On the contrary, the inhibition by blue light is stronger for *koy-1*: complete inhibition of mutant seeds is achieved at 56 µmol m^-2^ s^-1^ blue light whereas around 30% of WT seeds germinate under these conditions (Fig. 1E). These data indicate that germination control in *koy-1* mutant seeds is unresponsive to inhibition by red and far-red light, while the seeds perceive blue light.

Previously we showed that the light-sensitive germination of WT seeds might have evolved as a day-length sensing mechanism, allowing germination at short days in spring or autumn, and preventing it under long-day conditions (Merai et al., 2019). Applying the high light intensity of 450 µmol m^-2^ s^-1^ that inhibits even *koy-1* germination (Fig. 1A), we could show that the diurnal regulation was abolished in the mutant: *koy-1* seeds germinated irrespective of the day length regimes and were fully inhibited only in continuous light (Fig 1F).

### Genetic characterization of *koy-1*

Backcrosses of *koy-1* with the CYP wild type in both directions showed recessive inheritance of the light-unaffected germination, suggesting to map the causative mutation by searching for a homozygous polymorphism distinguishing mutant from wild-type siblings. We processed pools of 18 *koy-1* seedlings and 64 wild type progeny derived from the same M1 parent for DNA isolation and Illumina sequencing and mapped the reads to PacBio contigs of the CYP accession genome. We called single nucleotide polymorphisms (SNPs) and InDel variations in the *koy-1* pool, filtered against the control pool and asked for overlap with genic features. Thereby, we identified a 324 bp deletion in the promoter region of the heme oxygenase gene *AearHO1* (Fig. 2A,B), a gene closest to the Arabidopsis ortholog *AtHO1* according to phylogenetic and protein sequence analysis (Fig. 2C,D). The gene encodes a heme oxygenase, an enzyme involved in the biosynthesis of the phytochrome-associated chromophore, for which all enzymes and corresponding genes are known (Kohchi et al., 2001; Terry et al., 2002). In *Arabidopsis*, there are four heme oxygenases: group I with *AtHO1* (also called *HY1*), *AtHO3,* and *AtHO4*, and group II with *AtHO2* (Fig. 2C and Supplemental Fig. 2). The *Arabidopsis hy1/ho3/ho4* triple mutant shows severe symptoms with growth abnormalities, whereas milder effects in the single mutants indicate partial redundancy among the members of the group I (Emborg et al., 2006). Some *Brassicaceae* species have three heme oxygenases, while higher angiosperms often have only two heme oxygenase genes, corresponding to *HO1* and *HO2*, respectively (Fig. 2C and Supplemental Fig. S2). In *A. arabicum*, representing the earliest diverged sister group within the *Brassicaceae*, only two heme oxygenases could be identified with a protein Blast search, one of each HO group (Fig. 2C and Supplemental Fig. S2). The mutation in *koy-1* has removed the *AearHO1* promoter region between -90 and -414 bp upstream of the start codon, including the transcriptional start site identified by transcriptome analysis in the wild type (Fernandez-Pozo et al., 2021) and several binding sites for transcription factors (Fig. 2B,E). The expression level of *AearHO1* in seedlings of *koy-1* is reduced to 15% (Fig. 2F). The remaining transcripts are possibly initiated by the CAAT and TATA boxes between -414 and -500 bp upstream of the deletion (Fig. 2B,E). Genetic analysis in back-crossed progeny confirmed the co-segregation of the *AearHO1* promoter deletion with the ability to germinate in light, the phenotype of *koy-1* (Supplemental Table S3).

**Figure 2.**
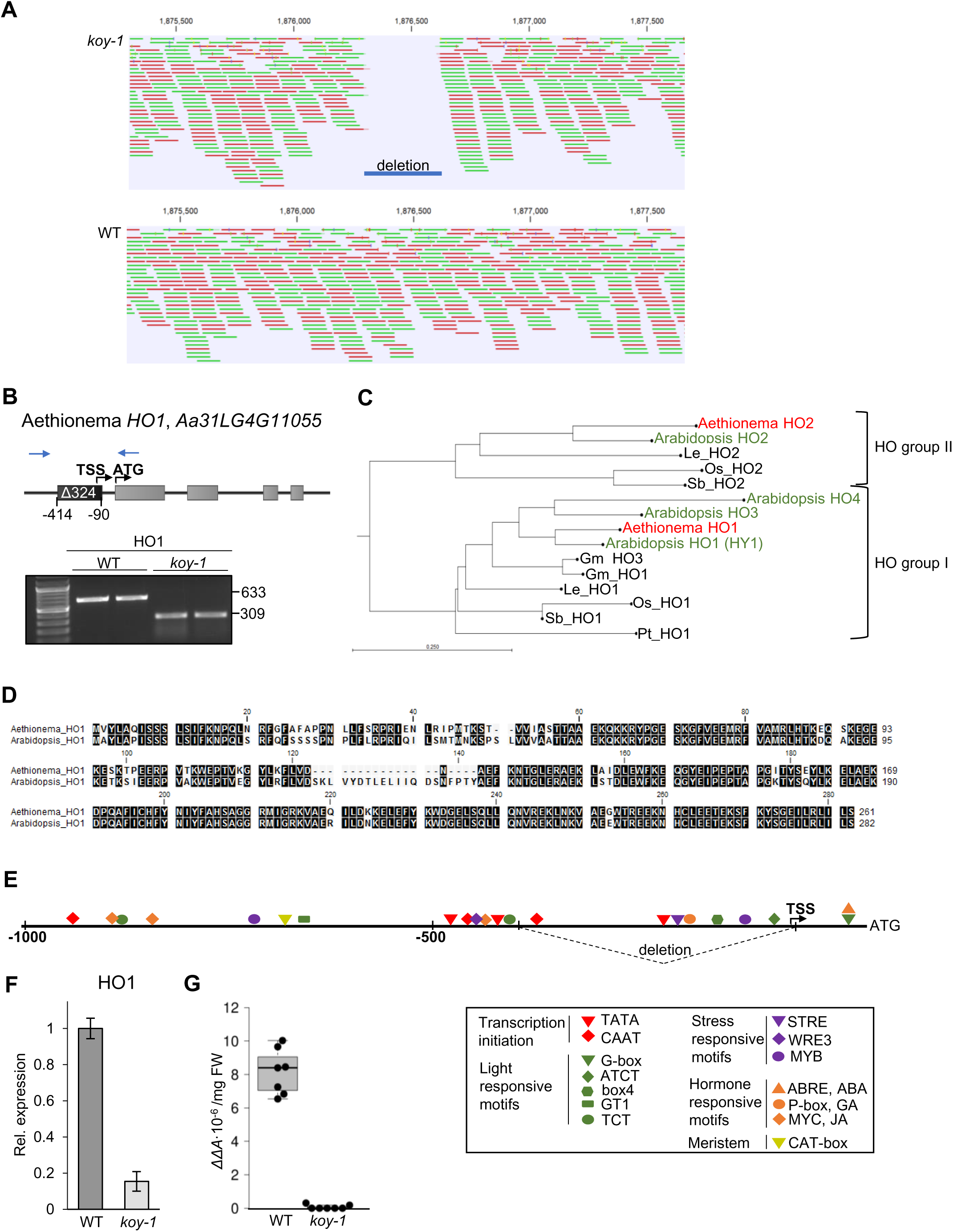
Identification of *koy-1* as mutation of the HEME OXYGENASE 1 gene. (A) Visualization of the mapped reads around the HO1 gene in the *koy-1* and WT pools. Green and red bars represent forward and reverse reads. (B) Simplified phylogenetic tree of genes of the heme oxygenase group. A complete version and a description are provided in Supplementary Figure S2 and Supplementary Table S5. (C) Schematic representation of the AearHO1 gene with the promoter deletion and its confirmation by PCR. (D) Protein alignment of Arabidopsis and Aethionema HO1. (E) Consensus motifs in the 1 kb region upstream of the ATG identified by PlantCARE. (F) Relative expression of *AearHO1* in dark- grown seedlings, normalized to the average WT level. Error bars indicate standard deviation of three biological replicates. (G) *In vivo* spectroscopy of photoreversible phytochromes from etiolated seedlings.

The role of HO1 in the biosynthesis of the phytochrome chromophore suggested that the strongly reduced expression (Fig. 2F) would reduce the amount of light-responsive phytochrome. Indeed, photoreversible phytochrome was hardly detectable in *koy-1*by in vivo spectroscopyin, in contrast to a phytochrome signal of >8 ΔΔA x 10^-6^/mg fresh weight in the wild type (Fig. 2G). In summary, the deletion at the *HO1* gene is limiting the amount of chromophore and photoreversible phytochrome and thereby plausibly the mutation in *koy-1* that is responsible for the altered light responsiveness of the seeds.

### Physiological characterization of *koy-1*

Lack of chromophore and active phytochromes is expected to affect light responses other than that on seed germination control. Therefore, we compared the hypocotyl elongation in darkness and under monochromatic light of different intensities. WT and *koy-1* mutant seedlings show the expected etiolated phenotype with elongated hypocotyls in darkness (Fig. 3A). The blue light suppressed the hypocotyl elongation to a similar extent in WT and *koy-1* seedlings (Fig. 3A,B). In contrast, in *koy-1*, the red and far-red light failed to inhibit hypocotyl elongation, indicating that the reception of red and far-red light and its downstream effects on seedling development are strongly affected (Fig. 3B-D).

**Figure 3.**
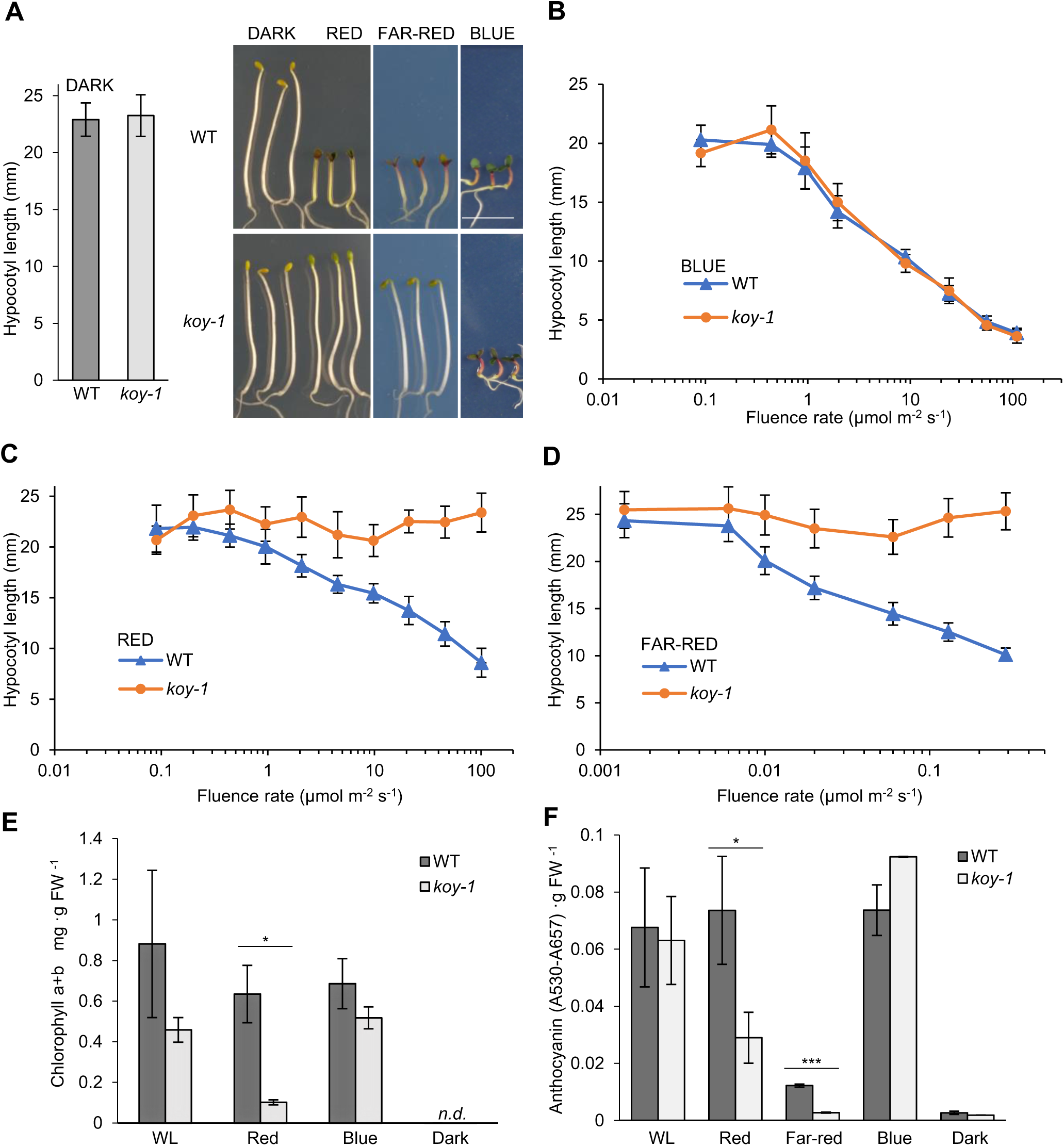
Photomorphogenic phenotype of WT and *koy-1* mutant seedlings. Hypocotyl length of WT and koy-1 mutant seedlings in darkness (A), blue (B), red (C), and far-red (D). Photos in (A) were taken from seedlings grown at the highest intensities applied in (B-D). Size bar represents 1 cm. Chlorophyll (E) and anthocyanin (F) content was measured in seedlings kept in darkness for germination induction followed by four days light exposure to 120 μmol m-2 s-1 white, 9 μmol m-2 s-1 red, 0.0008 μmol m-2 s-1 far-red, or 60 μmol m-2 s-1 blue light intensities. Error bars indicate standard deviations of three biological replicates. Asterisks indicate significant difference at * p<0.05, *** p<0.001 values tested by Student′s T-test.

To characterize further photomorphogenic traits, we determined chlorophyll and anthocyanin accumulation in seedlings grown under white, red, blue, far-red light or in darkness. The chlorophyll content was significantly lower in *koy-1* seedlings under red light, although the residual level might indicate a limited response (Fig. 3E). Anthocyanin accumulated in Aethionema seedlings under all light conditions, unlike in Arabidopsis, where red light does not induce this pigment (Neff and Chory, 1998). However, anthocyanin induction in *koy-1* mutants was significantly reduced under red and far-red light (Fig. 3F), further supporting the lack of perception of this light color by the mutant.

### Light-induced gene expression and hormonal changes in WT and *koy-1* seeds

To investigate the molecular basis of the light-inhibited seed germination in Aethionema, we examined the expression of key regulator genes under different light exposure and the amounts of hormones regulating germination. At imbibition, WT and *koy-1* seeds were illuminated for 24 h under 80 µmol m^-2^ s^-1^ red light or 1 µmol m^-2^ s^-1^ far-red light. Under these conditions, the germination of WT seeds is inhibited, while all *koy-1* seeds are germinated after three days. As the germination of *koy-1* seeds is hypersensitive to blue light, we chose 56 µmol m^-2^ s^-1^ blue light intensity, when WT seeds still retain ∼40% germination while the *koy-1* seeds are fully inhibited (Fig. 1E). Seeds imbibed in darkness were used as control, and harvested for sample preparation 24 h after imbibition like all others.

The expression level of *AearHO1* in wild-type and mutant seeds was independent from the light type but reduced to 10-13% in all *koy-1* mutant samples compared to the wild type (Fig. 4A), confirming the effect of the promoter deletion. *AearCHS*, encoding chalcone synthase and a commonly used light-regulated marker gene, was strongly induced by all light conditions in WT seeds, as expected. In *koy-1* seeds, only blue light induced *CHS* to a level comparable to the WT, confirming that the severe loss of phytochrome does not allow to respond to red and far-red light (Fig. 4B).

**Figure 4.**
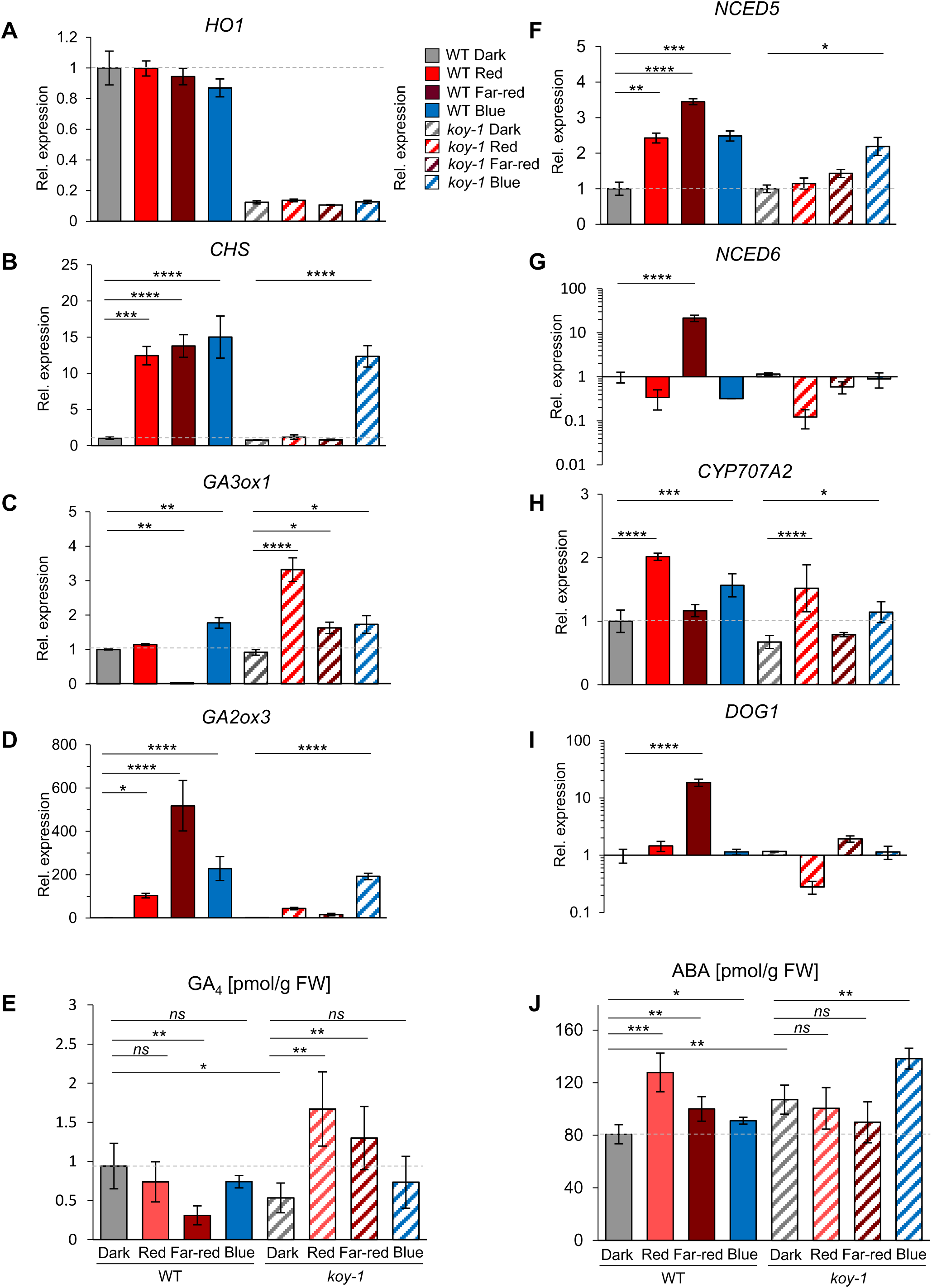
Light-induced gene expression and hormonal changes. (A-D and F-I) Quantitative RT–PCR for selected genes after 24 h light exposure with 80 µmol m-2 s-1 red light, 1 µmol m-2 s-1 far-red light, 56 µmol m-2 s-1 blue light, or kept in darkness. The gene expression levels in all light-exposed samples are presented as fold change relative to the average expression level in darkness of the same genotype. Expression in the dark in WT is set to 1 and indicated by the grey dashed lines. Asterisks indicate significant differences between the light- induced and dark expression level of a gene, based on the Tucay test: *p<0.05, **p<0.01, ***p<0.001, ****p<0.0001. Error bars represent standard deviation of three biological replicates. Hormone concentrations [in pmol g–1 fresh weight (FW)] of bioactive GA_4_ (E) and ABA (J) are shown for samples grown in dark, red, far-red, or blue light as in A-D and F-I. Levels in the dark in WT are indicated by grey dashed lines. Error bars represent standard deviation of five biological replicates. Asterisks indicate significant differences between the light-induced and dark values of one genotype or between the dark levels of WT and *koy-1*.

Seed germination is induced by shifting the hormonal balance towards higher gibberellin (GA) and lower abscisic acid (ABA) levels. The effect of light is exerted via transcriptional regulation of many genes involved GA and ABA metabolism, but in *Aethionema arabicum* often in the opposite direction compared to light-dependent germination in *Arabidopsis thaliana*, resulting a shifted hormonal balance towards ABA under white light (Mérai et al., 2019). Therefore, we tested the transcript levels of key regulatory genes and the hormonal levels in WT and *koy-1* mutant plants under red, far-red, or blue light. The expression of the GA biosynthetic enzyme *AearGA3ox1* is strongly reduced in far-red light in WT seeds but increased in *koy-1* seeds upon red and far-red treatment (Fig. 4C). Transcripts for the GA degradation enzyme *AearGA2ox3* increased significantly under red and far-red light in WT seeds only (Fig. 4D). Blue light significantly increased the *AearGA3ox1* and *AearGA2ox3* transcripts to similar level in WT and *koy-1* (Fig. 4C,D). In line with the transcriptional data, we detected a significant decrease of GA_4_ hormone level in WT seeds under far-red light. In contrast, GA_4_ is significantly increased in *koy-1* seeds under red and far-red light (Fig. 4E).

Among the ABA-regulating NCED genes encoding the 9-cis-epoxycarotenoid dioxygenases, which mediate a rate-limiting step of ABA synthesis, *AearNCED5* and *AearNCED6* were previously found to be induced by white light in the CYP accession of Aethionema (Mérai et al., 2019). The expression of *AearNCED5* in WT is increased in all light conditions, but in *koy-1* seeds only under blue light (Fig. 4F). The induction of *AearNCED6* is specific to far-red light in WT seeds but missing in the mutant (Fig. 4G). In contrast to the antipodal light regulation of *AearGA3ox1*, *AearGA2ox3*, *AearNCED5,* and *AearNCED6* between Aethionema and Arabidopsis seeds (Seo et al., 2006; Oh et al., 2007; Stawska and Oracz, 2015; Mérai et al., 2019), the ABA-deactivating enzyme *AearCYP707A2* is similarly induced by red or blue light, and reduced by far-red, in WT and *koy-1* mutant (Fig. 4H). Expression of *DELAY OF GERMINATION-1* (*DOG1*), an important regulator of seed dormancy via ABA synthesis, parallels that of NCED6 (Fig 4I). The ABA hormone levels were significantly increased in WT seeds in all light conditions, while only under blue light in mutant seeds (Fig. 4J). Moreover, the inhibition of *de novo* ABA synthesis by norflurazon completely rescued the WT seed germination both under red and far-red light, indicating a major role of ABA in the phytochrome-mediated germination inhibition (Supplemental Fig. S3). The addition of 100 µM GA_4+7_ also rescued the germination under red light, although the seeds needed longer time to overcome the inhibitory effect (Supplemental Fig. S3). Taken together, the transcriptional regulation of enzymes and the resulting amount and ratio between major hormonal determinants of germination are all concordant with the germination response of WT and *koy-1* mutant seeds to specific monochromatic light detected by phytochromes.

### Dual action of phytochromes on Aethionema seed germination

Our findings so far indicated that (1) the *koy-1* mutant lacks the red and far-red light-induced ABA accumulation and (2) the GA_4_ hormone level is not only unchanged in the mutant, but significantly increased under red and far-red. The latter seems to be in contrast to a lack of phytochrome activity. However, the residual (∼10%) expression of *AearHO1* might allow the production of small amounts of photoreversible phytochrome in the *koy-1* mutant, which is also suggested by reduced but measurable chlorophyll and anthocyanin amounts under red light. In *Arabidopsis*, the positive photoblastic germination can be induced by the very low fluence response (VLFR) through phyA, which requires as low as 0.1% of total phytochrome to be in the P_fr_ form, or by the low fluence response (LFR) through phyB (Botto et al., 1996; Shinomura et al., 1996). Dormant Arabidopsis seeds contain only phyB protein, while phyA is newly synthetized upon imbibition (Konomi et al., 1987; Shinomura et al., 1994; Shinomura et al., 1996). Shortly upon imbibition, phyB mediates the classical red/far-red photoreversible germination induction. Forty-eight hours later, the induction is photo-irreversible, as both red and far-red pulses trigger germination through phyA (Shinomura et al., 1996). Therefore, we aimed to elucidate which of the phytochrome action modes plays a role in the germination of Aethionema, which is strongly inhibited by light in a dosage-dependent way (Fig. 1) (Mérai et al., 2019). Knowing that the germination of Aethionema CYP wild-type seeds is neither inhibited by a 5 min red or far-red light pulse nor a 24 h red or far-red illumination (Mérai et al., 2019) clearly indicates that the inhibition is not a VLFR response. As both red and far-red light inhibits the germination, it is also unlikely that the inhibition is a low-fluence response (LFR), which is characterized by the red/far-red reversibility (Casal et al., 1998). Furthermore, seeds fully germinate after a far-red pulse, regardless if this is applied at 0, 3, 6 or 18 h after seed imbibition, while continuous far-red light strongly inhibits (Supplemental Fig. S4). This points to the third type of phytochrome response, high irradiance response (HIR), in which light pulses interrupted by dark cannot induce the same degree of response as continuous illumination, despite the same total fluence over time (Casal et al., 1998). To test whether the germination inhibition has HIR features, we compared WT seed germination either in constant far-red or red light or with hourly light pulses of 15 min followed by 45 min darkness. Importantly, the intermitted far-red or red light did not inhibit the germination, even at 3-4 orders of magnitude higher total fluence (Fig. 5A). This confirms that germination inhibition by light via phytochrome is an HIR response.

**Figure 5.**
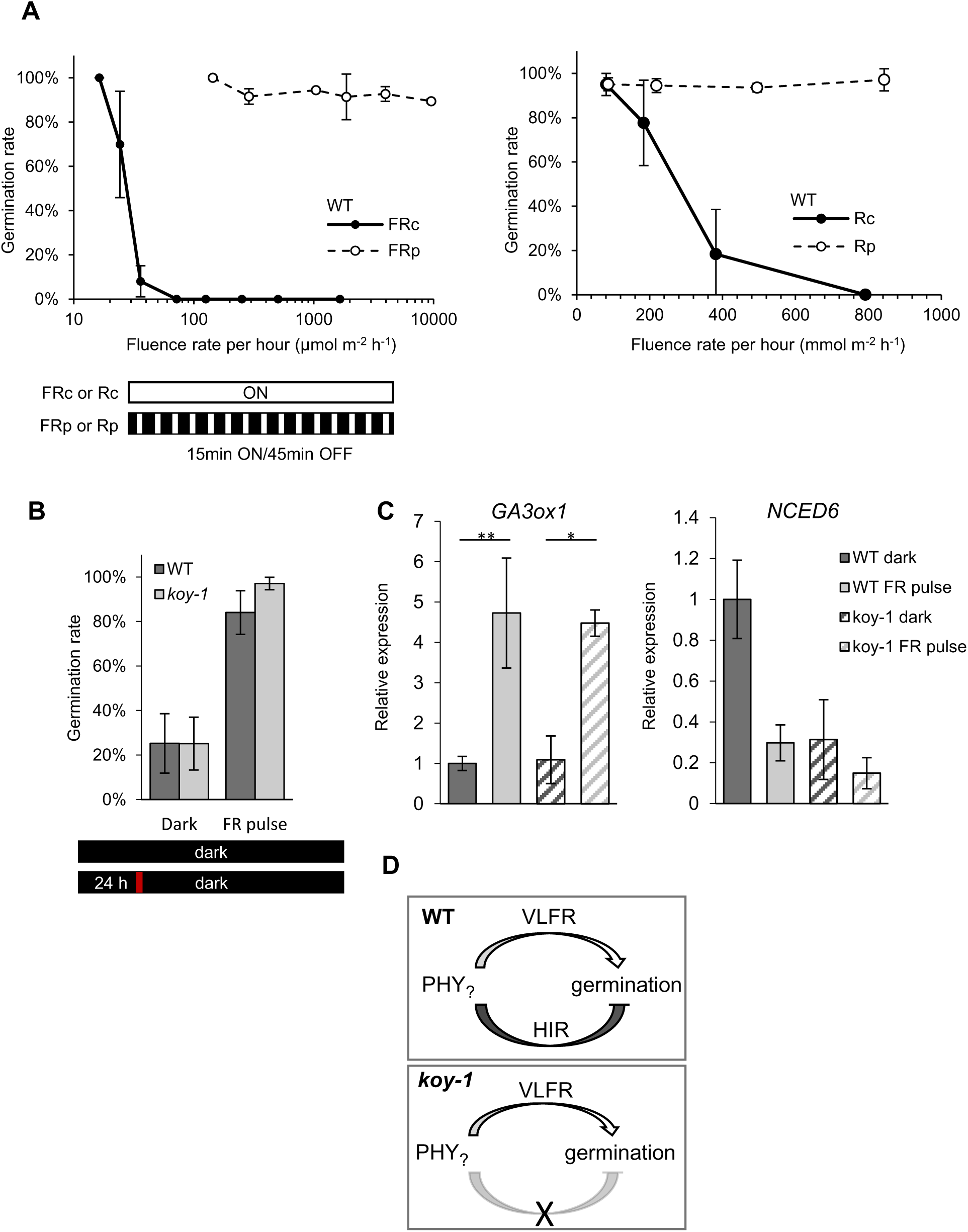
Antipodal effect on germination in high irradiance and very low fluence responses. (A) Germination rate of WT seeds under constant far-red or red light (FRc or Rc black line) or 15 min far-red or red pulses (FRp or Rp dashed line) intermitted with 45 min dark periods. (B) Germination rate of 8-week-old semi-dormant WT and *koy-1* seed batches in darkness with or without a 5 min far-red pulse 24 h after imbibition. (C) Quantitative RT-PCR of *GA3ox1* and *NCED6* from seeds described in (B). Samples were collected 30 h after imbibition; 6 h after the far-red pulse. Asterisks indicate significant differences based on the Tucay test: *p<0.05, ** p<0.01. Error bars represent standard deviation of three biological replicates. (D) Schematic model of dual action of phytochromes. The germination induction by very low fluence response (VLFR) is present in both WT and *koy-1* mutant seeds, while the germination-inhibiting high irradiance response (HIR) only operates in WT seeds.

The increasing GA_4_ hormone level in the *koy-1* mutant under red and far-red light (Fig. 4E,J) indicated that, under certain circumstances, light exposure might support rather than inhibit seed germination. As the residual expression of *AearHO1* in *koy-1,* as discussed, is possibly sufficient for a VLFR response, we hypothesized that VLFR might induce germination, similar to *Arabidopsis*. As non-dormant Aethionema seeds germinate in darkness without any light, we tested 8-week-old semi-dormant WT and *koy-1* seed batches with a limited germination rate of around 25% in darkness, allowing us to see a positive effect of light exposure. We performed germination assays in complete darkness, with or without a 5 min far-red pulse 24 h after imbibition (Fig. 5B). Indeed, the light pulse increased the germination to 84% and 97% in WT and *koy-1* seeds, respectively, indicating that the short light exposure triggers VLFR in both lines (Fig. 5B). Concomitantly, seeds that received the 5 min far-red pulse had enhanced expression of *AearGA3ox1* and reduced expression of *AearNCED6* (Fig. 5C). All data taken together indicate that phytochromes have two, antipodal effects on seed germination in the CYP accession of *A. arabicum*: continuous red and far-red light strongly inhibits germination in a dosage-dependent manner, while a short light pulse can induce germination (Fig. 5D). Mature seeds that have lost the primary dormancy germinate in darkness without needing any light, but the seeds retain the innate module of VLFR induction prior to full maturation.

### Fruit and seed formation in WT and *koy-1* plants

The reduced level of HO1 and photoreversible phytochrome let us expect additional effects besides altered regulation of seed germination, seedling development and pigment synthesis. Double and triple mutants of the *HO1* gene family in Arabidopsis display severe growth abnormalities, small chlorotic leaves, and early flowering (Emborg et al., 2006). Aethionema *koy*-*1* mutant plants grow well but resemble a shade-avoidance response (Keuskamp et al., 2010), including a tall, elongated stem, long hypocotyls and internodes, and pale green color (Fig. 6A,B). Unlike in many species, we did not observe earlier flowering, longer petioles, or modified branching patterns (Fig. 6B). A specific feature of *A. arabicum* is the formation of heteromorphic fruits: the same individual plant generates both, indehiscent (IND) and dehiscent (DEH) fruit morphs enclosing either a single or up to six seeds, respectively (Fig. 7A) and (Lenser et al., 2016). Both fruit types can appear also with aborted seeds (Fig. 7A). Previous experiments mainly made with the Aethionema accession originating from Turkey had shown that the ratio of indehiscent versus dehiscent fruit morphs is a plastic phenotype influenced by various factors (Lenser et al., 2016; Bhattacharya et al., 2019), in response to auxin treatment, defoliation and shading. Given the shade-avoidance phenotype of *koy-1* plants, we tested if fruit and seed production or the fruit morph ratio in the mutant would be different compared to the CYP wild type. Plants were grown under the same spectral, diurnal, and temperature conditions but either under 300 µmol m^-2^ s^-1^ light intensity, the standard laboratory growth condition for *A. arabicum*, or at lower, suboptimal 200 µmol m^-2^ s^-1^ light intensity. The wild-type plants were highly sensitive to the light condition: although they produced more fruits under the limiting light, these had less viable seeds than under regular light (Fig. 7B,C) and lower seed numbers in the seeded DEH fruits (Fig. 7D). In contrast, *koy-1* plants tolerated the lower light intensity much better, with even slightly higher fruit numbers, comparable total seed numbers, and only minor reduction of seeded DEH fruits (Fig. 7B-D).

**Figure 6.**
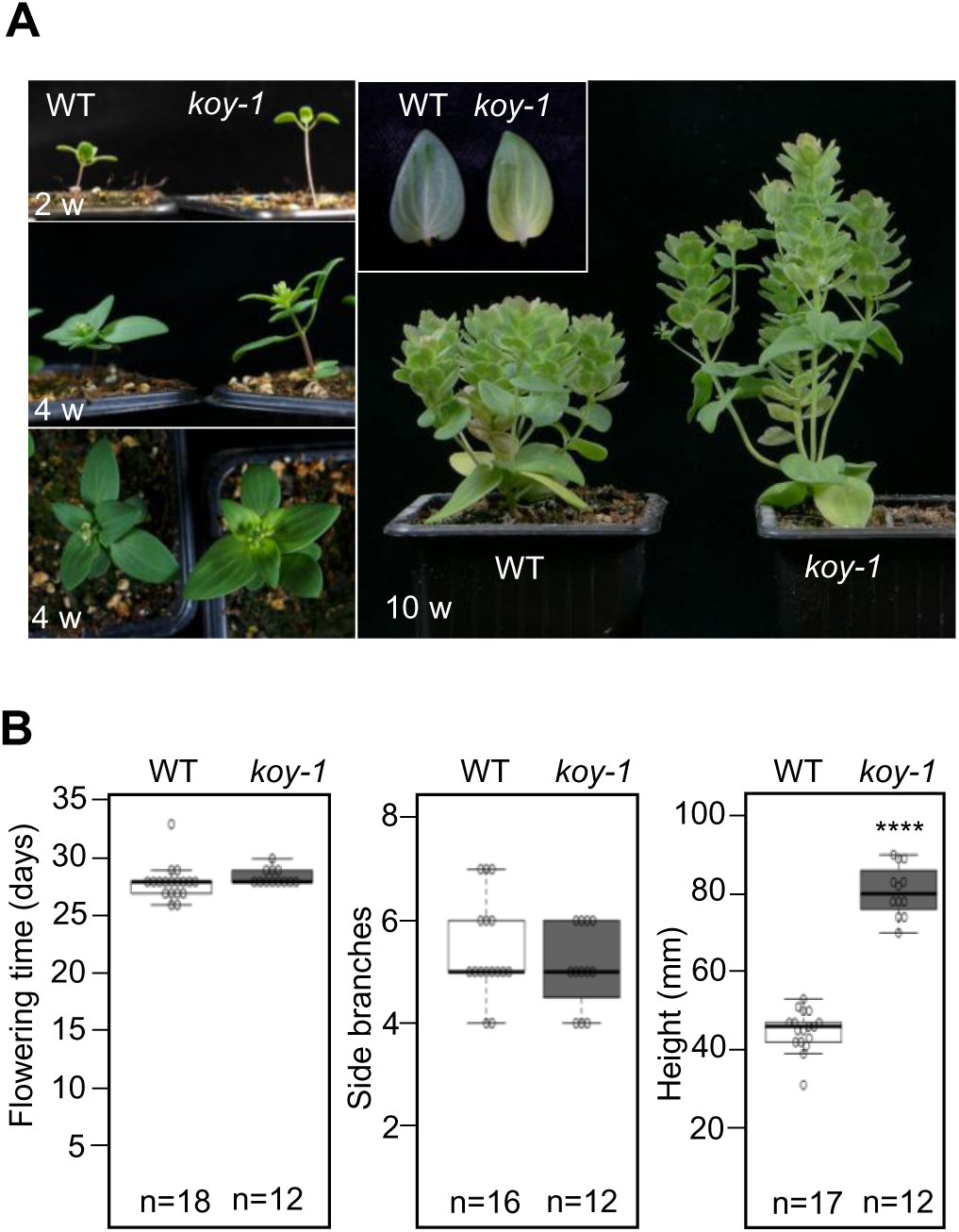
Phenotype of WT and *koy-1* mutant plants. (A) Photos of two-, four- or ten-week-old plants. (B) Flowering time, number of primary side branches, and plant height was recorded for 12-18 plants per box plot. Asterisks indicate significant difference at **** p<0.0001 tested with Student′s T-test.

**Figure 7.**
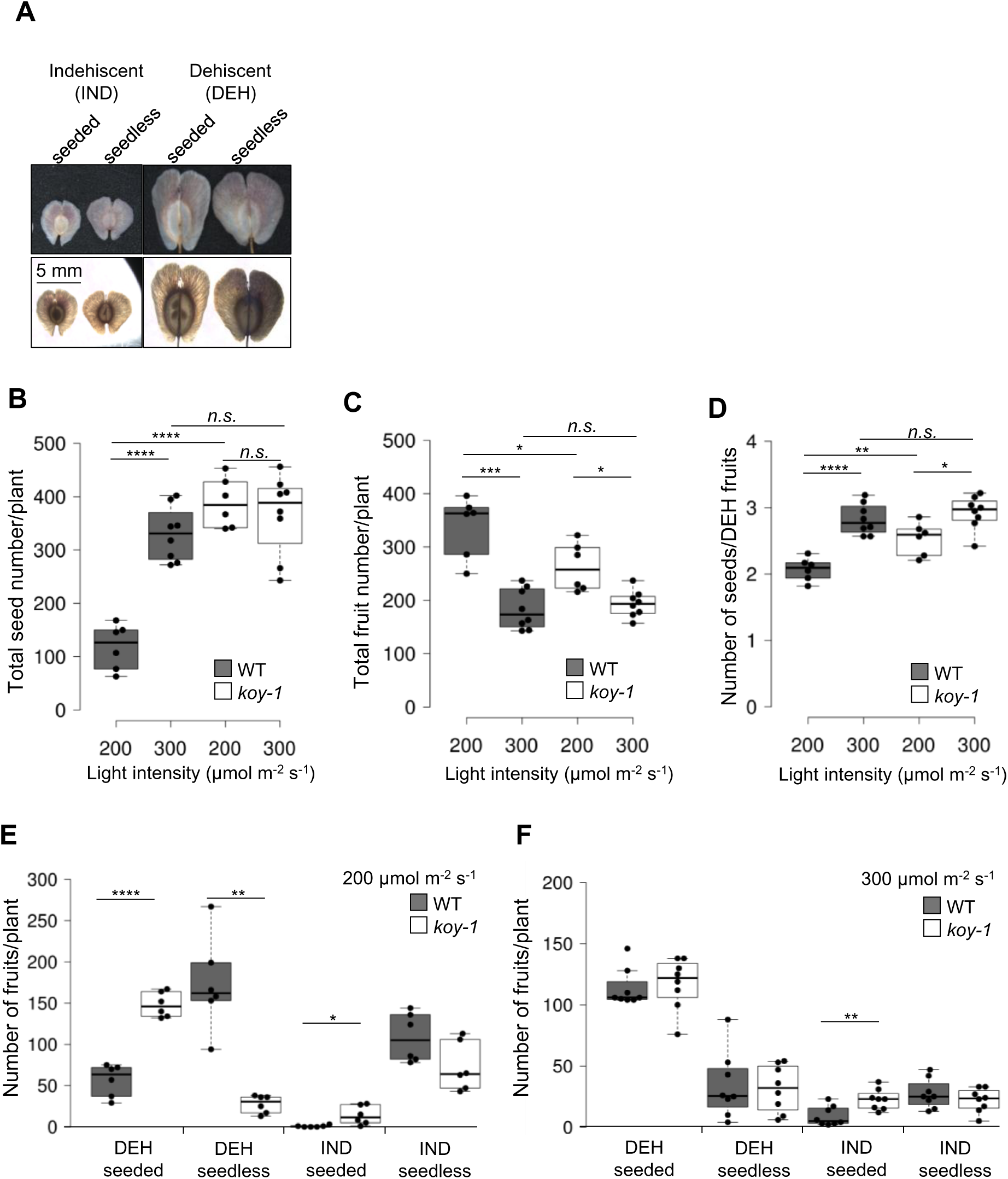
Fruit and seed production and fruit morph ration in WT and *koy-1* mutant plants. (A) Appearance of indehiscent (IND) and dehiscent (DEH) fruit types with or without seeds. Note that the seed number in the DEH fruit type varies from one to six. (B-F) Seed and fruit numbers of fully matured plants kept under 200 or 300 μmol m-2 s-1 white light from germination on. Center lines show the medians; box limits indicate the 25th and 75th percentiles, whiskers extend 1.5 times the interquartile range from the 25th and 75th percentiles. N = 6 or 8, for 200 and 300 μmol m-2 s-1 intensities, respectively. Asterisks indicate significant differences with *p<0.05, **p<0.01, ***p<0.001, ****p<0.0001 or n.s. as no significant difference, tested with Student′s T-test. Error bars represent the standard deviation from three biological replicates.

While DEH fruit morphs shed the valves upon maturity so that seeds fall off and remain close to the mother plant, IND fruit morphs keep enclosing a single seed and have a winglet-like structure with high dispersal ability (Fig. 7A) (Lenser et al., 2016; Bhattacharya et al., 2019). In the TUR accession, seeds within IND fruits have stronger dormancy (Lenser et al., 2016; Arshad et al., 2019; Bhattacharya et al., 2019). Therefore, the IND fruit type provides spatio-temporal flexibility for colonization compared to seeds originating from DEH fruits (Bhattacharya et al., 2019). Interestingly, while the WT plants did not produce seeded IND fruits under low light and only a few under 300 µmol m^-2^ s^-1^, *koy-1* plants have significantly more seeded IND fruits at both light intensities (Fig. 7E,F). At 300 µmol m^-2^ s^-1^ , the number of seeded IND fruits is the only significantly different parameter between *koy-1* and WT, while the number of DEH fruits and seeds per fruit are similar (Fig. 7D,F). These results indicate that phytochromes affect the tolerance of lower light intensity and its effect on fruit and seed production in *Aethionema arabicum*.

## Discussion

Although negative photoblasty is a common feature with multiple evolutionary origin, it contrasts with current textbook knowledge, and its molecular basis is only scarcely understood. Therefore, the natural variation of the trait in a *Brassicaceae* species, closely related to the classical model *Arabidopsis thaliana,* and the highly improved gene annotation in a relatively small genome (Fernandez-Pozo et al., 2021) recommend *A. arabicum* as an attractive novel model to investigate the complex regulation of seed germination in response to light. We created the first mutant collection for *A. arabicum* to perform a forward genetic screen for lack of white light-inhibited germination. In this study we present the characterization of the first mutant, harboring a promoter deletion in the gene *HEME OXYGENASE 1*, encoding a key enzyme for chromophore biosynthesis. The difference between this mutant and the parental WT allowed us to explain differences in the light response compared to Arabidopsis and additional phenotypic changes mediated by light reception via phytochromes.

### Phytochrome action in WT and *koy-1* seeds

As there is only one member of the heme oxygenase group I in Aethionema, it was plausible that the deletion in the promoter of the *AearHO1* gene in the *koy-1* mutant resulted in a strongly reduced amount of photoconvertible phytochromes, confirmed by the resulting effects on hypocotyl growth, seed germination, the lack of *AearCHS* induction under red and far-red light, and the lack of anthocyanin accumulation under far-red light. Moreover, the *koy-1* plant phenotype resembles that of Arabidopsis *phyB* null mutants. However, residual chlorophyll and anthocyanin accumulation under red light indicated that a minor portion of phytochromes can still be activated in *koy*-1.

We demonstrated that the phytochromes play a dual, antipodal role in the germination of Aethionema (CYP) seeds: they inhibit the germination in an HIR action mode but induce the germination through the VLFR. In WT seeds, increasing light intensity and duration shift the positive effect of VLFR towards the inhibition by HIR. Importantly, the residual amount of photoreversible phytochrome in *koy-1* is sufficient for the VLFR induction, but not for the inhibition through the HIR. The *koy-1* mutant allows the genetic dissection of the two response types. For example, VLFR and HIR both control the GA_4_ level, as the *AearGA3ox1* expression and the GA_4_ hormone accumulation show contrasting response between WT and mutant. In contrast, the degradation of GA_4_ through GA3ox2 is only an HIR response, as there was no significant induction in *koy-1*. The ABA synthesis through NCED5 and NCED6 is an HIR response, while the ABA degradation via CYP707A2 might be mainly due to VLFR, as the expression profile is similar in WT and *koy-1* seeds. Similar phenomena were found in *Datura ferox* seeds, where the far-red HIR antagonized the VLFR induction of the *DfGA3ox* gene (Arana et al., 2007).

As VLFR and HIR upon far-red exposure are mediated by the PhyA photoreceptor, it is very likely that PhyA has a major role in the control of seed germination in Aethionema. However, the presented data do not exclude the role of other phytochromes and the LFR type in the process, particularly in the dosage-dependent red-light inhibition. The PhyA-mediated germination inhibition by far-red HIR is well known from tomato (Appenroth et al., 2006; Auge et al., 2009). A limited number of studies reports germination inhibition by continuous red light in Californian poppy seeds and *Phacelia tanacetifolia,* or by red light regimes in *Bromus sterilis*, but the specific responsible phytochrome is unknown (Schulz and Klein, 1963; Goldthwaite et al., 1971; Hilton, 1982). The mutant population established in *Aethionema arabicum* could likely provide a valuable resource to screen for individual phytochrome mutants to study their role in the inhibition of seed germination.

### Two distinct pathways for light inhibited germination in Aethionema

Loss of seed germination inhibition in the *koy-1* mutant in red and far-red light, but its persistence under blue light, indicates the existence of at least two pathways for the light response in Aethionema: a phytochrome-mediated red and far-red inhibition, and a blue light-induced, phytochrome-independent pathway. This also explains the inhibition of *koy-1* seeds under high intensity white light. The role of blue light in seed dormancy and germination depends strongly on the taxon. In monocots, blue light inhibition of seed germination is reported for barley, ryegrass, and *Brachypodium* (Goggin et al., 2008; Gubler et al., 2008; Barrero et al., 2012). RNAi application in barley revealed that this is mediated through the CRY1 photoreceptor, which induces the ABA synthesis (Barrero et al., 2014). In dicots, inhibitory effects of blue light on seed germination are described for watermelon, *Laportea bulbifera,* and *Trifolium subterraneum* (Tanno, 1983; Thanos and Mitrakos, 1992; Costa et al., 2016). The lack of genetic tools in these species so far does not allow concluding which photoreceptor/s mediate the inhibition. In *Arabidopsis*, blue light has a positive effect on dormancy alleviation through the PHYB photoreceptor (Stawska and Oracz, 2019). Interestingly, our results indicate a phytochrome-independent blue light inhibition in a dicot plant.

The inhibition of germination under blue light, stronger in the *koy-1* mutant than in wild type, might indicate an interplay between the two pathways. Interactions between the phytochromes and the blue light photoreceptor CRY1 have been demonstrated in many aspects, including hypocotyl growth, cotyledon unfolding and expansion, chlorophyll and anthocyanin accumulation (Casal and Boccalandro, 1995; Ahmad and Cashmore, 1997; Neff and Chory, 1998). However, in contrast to *Arabidopsis* phytochrome mutants (Neff and Chory, 1998), we did not observe differences in hypocotyl length, chlorophyll or anthocyanin content between WT and *koy-1* seedlings under blue light. Alternatively, the stronger inhibition by blue light in *koy-1* might be a consequence of lower GA_4_ and higher ABA levels in the seed before imbibition. Even if the ABA induction upon blue light is equal in WT and *koy-1* seeds, it could result in higher absolute ABA levels if the initial ABA level was already higher in the mutant. This explanation is supported by the similar expression patterns of key regulator genes under blue light in both genotypes and the higher ABA level in *koy-1* in the dark samples.

### The ecological role of phytochrome-mediated germination control

Several data presented before and here suggest that the ecological role of light-inhibited germination is likely a day-length sensing mechanism to ensure the appropriate timing of germination in the original habitat of the Cyprus accession (Mérai et al., 2019). Besides moisture, a temperature range around 14°C is necessary for *A. arabicum* seed germination (Arshad et al., 2019), but these conditions can occur in other seasons. Combining these requirements with that for short day length restricts the germination period to a narrow time-window in early spring. Consequently, the plants finish their life cycle of ∼4 months before the dry and warm summer (Bhattacharya et al., 2019; Mérai et al., 2019). Other examples of light-inhibited germination can be found among desert or Mediterranean maritime plants, which are also challenged by drought, heat, and high light exposure (Thanos et al., 1991; Lai et al., 2016; Carta et al., 2017). We demonstrated that the diurnal regulation of germination is mediated by phytochromes, as *koy-1* mutant germinates under all diurnal regimes. On the other hand, *A. arabicum* accessions from Turkey and the closely related species *A. heterocarpum* originating from Israel have light-neutral seeds that germinate equally well in darkness and light; therefore, their germination does not depend on the day length (Merai et al., 2019). Why other closely related Aethionema accessions from similar climate conditions did not acquire, or have lost, the potential advantages of the day-length sensing mechanism, remains an open question. The evolution of germination strategies is not independent of other traits, for example the formation of soil seed bank, alternative risk-reducing traits like stress-tolerant morphology, seed size, or bet-hedging strategies (Venable and Brown, 1988; Saatkamp et al., 2019). *A. arabicum* is a fascinating model for a double bet-hedging strategy with a spatial and temporal dispersal dimorphism. The multi-seeded dehiscent fruits release seeds with low dormancy close to the parental plant, while the single-seeded indehiscent fruits with winglet-like pericarps are dispersed by wind or water currents and have pronounced dormancy due to much more ABA compared to the DEH seeds (Lenser et al., 2016; Arshad et al., 2019). In most other dimorphic systems described, high dispersal probability is combined with less dormancy, and *vice versa* (Venable and Lawlor, 1980; Baskin et al., 2014). The opposite correlation, high dormancy of far distributed seeds in Aethionema, make the appearance and amount of IND fruits a key factor for the bet-hedging strategy (Arshad et al., 2019). Interestingly, the ratio between the two fruit morphs strongly varies among the accessions. In field data collected in the original habitat of the TUR accession, the IND:DEH ratio was between 0.52-0.58:1 (Bhattacharya et al., 2019); data for CYP in its original habitat are not available. In controlled conditions in growth chambers, the ratio in TUR plants was shifted towards more IND fruit (2.5:1), whereas the CYP accession produced only a few IND fruits (0.08:1, IND:DEH). *koy-1* mutants produce 2.5 times more IND fruits, resulting in a significant increase in the absolute number of IND fruits and a higher (0.19:1) IND:DEH fruit morph ratio(Supplemental Fig. S5). Whether and how the loss of light sensitivity of the *koy-1* mutant is functionally connected with the appearance of more IND fruits remains to be explored, but it is not unlikely, as the TUR accession with fully light-insensitive seeds has substantially more IND fruits. Such a connection might reflect alternative strategies: Aethionema accessions originating from semi-arid habitats have either a precise timing of germination controlled by multiple factors including day length sensitivity, or a bet-hedging strategy with more IND fruits and spatiotemporal dispersal at the cost of the loss of light sensitivity and seasonal control (Supplemental Fig. S5B). Future research with more accessions can address if there is a negative correlation between light sensitivity and dimorphic dispersal of seeds. Independent of this potentially adaptive aspect, the already available range of natural diversity within the Aethionemeae and growing genetic and genomic resources in one of its members open avenues to study several traits of ecological and physiological relevance that are not accessible in other model plants.

## Supplemental information

**Supplemental Figure S1** Temperature-dependent seed germination of WT and koy-1 mutant seeds.

**Supplemental Figure S2** Phylogram of heme oxygenases in seed plants, related to Figure 3B.

**Supplemental Figure S3** Germination rescue with norflurazon and gibberellin.

Percentage of germination over time is shown under 110 µmol m^-2^ h^-1^ red (C) and 0.3 µmol m^-2^ h^-1^ far-red (B) light with the addition of 100 µM GA_4+7_ (GA) or 50 µM norflurazon (NOR) or 0.01% DMSO as a control. Dark (A) is shown as control without treatment. Error bars represent standard deviation of three independent replicates.

**Supplemental Figure S4** The effect of far-red pulses on seed germination during imbibition.

Germination of WT and *koy-1* seeds after 6 days in continuous darkness (cDark), under continuous far-red (cFR) or in darkness after a 5 min far-red pulse upon imbibition, or 3, 6, 18 hours after imbibition. Error bars represent standard deviation of three independent replicates.

**Supplemental Figure S5** Propagation strategies in *A. arabicum*.

(A) Fruit-morph ratio in two wild-type accessions and the *koy-1* mutant. Asterisks indicate significant difference at ** p<0.01, *** p<0.001 values tested by Student′s T-test. (B) Hypothetical model of alternative strategies in *Aethionema* wild-type accessions and the *koy-1* mutant. -* indicates that the *koy-1* seeds exhibit hyposensitivity to white light but have lost diurnal regulation.

**Supplemental Table S1** List of Aethionema accession numbers used for this study.

**Supplemental Table S2** List of primers used for quantitative RT-PCR analysis.

**Supplemental Table S3** Co-segregation of light germination phenotype under white light with the promoter deletion identified in koy-1.

**Supplemental Table S4** Co-segregation of long hypocotyl phenotype under far-red with the promoter deletion identified in koy-1.

**Supplemental Dataset 1** Protein sequences used for phylogenetic tree, related to Supplementary Figure 2.

## Acknowledgements

We thank to Eric M. Schranz for providing Aethionema seed stocks. We also thank the staff of the Vienna BioCenter Core Facilities GmbH (VBCF), a member of Vienna BioCenter (VBC), Austria, especially the Plant Sciences Facility for growth of the plants and the wavelength-specific light experiments, the Next Generation Sequencing Facility for generating the PacBio and whole-genome sequencing data, the Molecular Biology Unit for providing multiple reagents, and the Vienna Covid-19 Detection Initiative (VCDI) for generating a safe work environment during the pandemic. We thank Nicole Lettner, Marie Vitaskova and Magdalena Vlckova for excellent technical support. SK was supported by the Refugee Support Program of the Austrian Academy of Sciences. The work was funded by the Austrian Science Fund (FWF) to ZM (FWF I3979-B25). It was additionally supported by the European Regional Development Fund Project ‘Centre for Experimental Plant Biology’ (CZ.02.1.01/0.0/0.0/16_019/0000738) to D.T. and by the Czech Science Foundation to M.S. (grant No. GA21-07661S).

## Author contributions

ZM and OMS planned and designed the research; ZM, FX, AM, CK, FA, SK, DT, and VT performed the experiments; ZM, FX, LMSJ, KL, CK, DT, VT, MS, and interpreted the data; ZM and OMS wrote the paper. All authors approved the submitted version.

## References

Ahmad, M., and Cashmore, A.R. (1997). The blue-light receptor cryptochrome 1 shows functional dependence on phytochrome A or phytochrome B in Arabidopsis thaliana. Plant J 11, 421–427.

Appenroth, K.J., Lenk, G., Goldau, L., and Sharma, R. (2006). Tomato seed germination: regulation of different response modes by phytochrome B2 and phytochrome A. Plant Cell Environ 29, 701–709.

Arana, M.V., Burgin, M.J., de Miguel, L.C., and Sánchez, R.A. (2007). The very-low-fluence and high-irradiance responses of the phytochromes have antagonistic effects on germination, mannan-degrading activities, and DfGA3ox transcript levels in Datura ferox seeds. J Exp Bot 58, 3997–4004.

Arshad, W., Sperber, K., Steinbrecher, T., Nichols, B., Jansen, V.A.A., Leubner-Metzger, G., and Mummenhoff, K. (2019). Dispersal biophysics and adaptive significance of dimorphic diaspores in the annual Aethionema arabicum (Brassicaceae). New Phytol 221, 1434–1446.

Auge, G.A., Perelman, S., Crocco, C.D., Sánchez, R.A., and Botto, J.F. (2009). Gene expression analysis of light-modulated germination in tomato seeds. New Phytol 183, 301–314.

Barragan, C.A., Collenberg, M., Schwab, R., Kerstens, M., Bezrukov, I., Bemm, F., Požárová, D., Kolář, F., and Weigel, D. (2021). Homozygosity at its limit: inbreeding depression in wild Arabidopsis arenosa populations. bioRxiv 2021.01.24.427284.

Barrero, J.M., Downie, A.B., Xu, Q., and Gubler, F. (2014). A role for barley CRYPTOCHROME1 in light regulation of grain dormancy and germination. Plant Cell 26, 1094–1104.

Barrero, J.M., Jacobsen, J.V., Talbot, M.J., White, R.G., Swain, S.M., Garvin, D.F., and Gubler, F. (2012). Grain dormancy and light quality effects on germination in the model grass *Brachypodium distachyon*. New Phytol 193, 376–386.

Baskin, J.M., Baskin, C.C., Tan, D.Y., and Wang, L. (2014). Diaspore dispersal ability and degree of dormancy in heteromorphic species of cold deserts of northwest China: a review. Perspectives in Plant Ecology, Evolution and Systematics 16, 93–99.

Bhattacharya, S., Sperber, K., Özüdoğru, B., Leubner-Metzger, G., and Mummenhoff, K. (2019). Naturally-primed life strategy plasticity of dimorphic Aethionema arabicum facilitates optimal habitat colonization. Sci Rep 9, 16108.

Botha, F.C., and Small, J.G.C. (1988). The germination response of the negatively photoblastic seeds of Citrullus lanatus to light of different spectral compositions. Journal of Plant Physiology 132, 750–753.

Botto, J.F., Sanchez, R.A., Whitelam, G.C., and Casal, J.J. (1996). Phytochrome A mediates the promotion of seed germination by very low fluences of light and canopy shade light in Arabidopsis. Plant Physiol 110, 439–444.

Brown, S.B., Houghton, J.D., and Vernon, D.I. (1990). Biosynthesis of phycobilins. Formation of the chromophore of phytochrome, phycocyanin and phycoerythrin. J Photochem Photobiol B 5, 3–23.

Carta, A., Skourti, E., Mattana, E., Vandelook, F., and Thanos, C.A. (2017). Photoinhibition of seed germination: occurrence, ecology and phylogeny. Seed Science Research 27, 131–153.

Casal, J.J., and Boccalandro, H. (1995). Co-action between phytochrome B and HY4 in Arabidopsis thaliana. Planta 197, 213–218.

Casal, J.J., and Sanchez, R.A. (1998). Phytochromes and seed germination. Seed Science Research 8, 3.

Casal, J.J., Sanchez, R.A., and Botto, J.F. (1998). Modes of action of phytochromes. Journal of Experimental Botany 49, 127–138.

Chen, M., MacGregor, D.R., Dave, A., Florance, H., Moore, K., Paszkiewicz, K., Smirnoff, N., Graham, I.A., and Penfield, S. (2014). Maternal temperature history activates Flowering Locus T in fruits to control progeny dormancy according to time of year. Proc Natl Acad Sci U S A 111, 18787–18792.

Chory, J., Peto, C., Feinbaum, R., Pratt, L., and Ausubel, F. (1989). Arabidopsis thaliana mutant that develops as a light-grown plant in the absence of light. Cell 58, 991–999.

Christie, J.M., Blackwood, L., Petersen, J., and Sullivan, S. (2015). Plant flavoprotein photoreceptors. Plant Cell Physiol 56, 401–413.

Costa, A., Soveral Dias, A., Grenho, M.G., and Silva Dias, L. (2016). Effects of dark or of red, blue or white light on germination of subterranean clover seeds. Emirates Journal of Food and Agriculture 28, 853–864.

Dehesh, K., Franci, C., Parks, B.M., Seeley, K.A., Short, T.W., Tepperman, J.M., and Quail, P.H. (1993). Arabidopsis HY8 locus encodes phytochrome A. Plant Cell 5, 1081–1088.

Emborg, T.J., Walker, J.M., Noh, B., and Vierstra, R.D. (2006). Multiple heme oxygenase family members contribute to the biosynthesis of the phytochrome chromophore in Arabidopsis. Plant Physiol 140, 856–868.

Fernandez-Pozo, N., Metz, T., Chandler, J.O., Gramzow, L., Mérai, Z., Maumus, F., Mittelsten Scheid, O., Theißen, G., Schranz, M.E., Leubner-Metzger, G., and Rensing, S.A. (2021). *Aethionema arabicum* genome annotation using PacBio full-length transcripts provides a valuable resource for seed dormancy and Brassicaceae evolution research. Plant J 106, 275–293.

Finch-Savage, W.E., and Leubner-Metzger, G. (2006). Seed dormancy and the control of germination. New Phytologist 171, 501–523.

Furuya, M., and Schäfer, E. (1996). Photoperception and signallingof induction reactions by different phytochromes. Trends in Plant Science 1, 301–307.

Goggin, D.E., Steadman, K.J., and Powles, S.B. (2008). Green and blue light photoreceptors are involved in maintenance of dormancy in imbibed annual ryegrass (*Lolium rigidum*) seeds. New Phytol 180, 81–89.

Goldthwaite, J.J., Bristol, J.C., Gentile, A.C., and Klein, R.M. (1971). Light-suppressed germination of California poppy seed. Canadian Journal of Botany 49, 1655–1659.

Gubler, F., Hughes, T., Waterhouse, P., and Jacobsen, J. (2008). Regulation of dormancy in barley by blue light and after-ripening: effects on abscisic acid and gibberellin metabolism. Plant Physiol 147, 886–896.

Gyula, N., Schafer, E., and Nagy, F. (2003). Light perception and signalling in higher plants. Current Opinion in Plant Biology 6, 446–452.

Haudry, A., Platts, A.E., Vello, E., Hoen, D.R., Leclercq, M., Williamson, R.J., Forczek, E., Joly-Lopez, Z., Steffen, J.G., Hazzouri, K.M., Dewar, K., Stinchcombe, J.R., Schoen, D.J., Wang, X.W., Schmutz, J., Town, C.D., Edger, P.P., Pires, J.C., Schumaker, K.S., Jarvis, D.E., Mandakova, T., Lysak, M.A., van den Bergh, E., Schranz, M.E., Harrison, P.M., Moses, A.M., Bureau, T.E., Wright, S.I., and Blanchette, M. (2013). An atlas of over 90,000 conserved noncoding sequences provides insight into crucifer regulatory regions. Nature Genetics 45, 891–U228.

Hilton, J.R. (1982). An unusual effect of the far-red absorbing form of phytochrome: Photoinhibition of seed germination in *Bromus sterilis* L. Planta 155, 524–528.

Hofmeister, B.T., Denkena, J., Colomé-Tatché, M., Shahryary, Y., Hazarika, R., Grimwood, J., Mamidi, S., Jenkins, J., Grabowski, P.P., Sreedasyam, A., Shu, S., Barry, K., Lail, K., Adam, C., Lipzen, A., Sorek, R., Kudrna, D., Talag, J., Wing, R., Hall, D.W., Jacobsen, D., Tuskan, G.A., Schmutz, J., Johannes, F., and Schmitz, R.J. (2020). A genome assembly and the somatic genetic and epigenetic mutation rate in a wild long-lived perennial *Populus trichocarpa*. Genome Biol 21, 259.

Holm, M., Ma, L.G., Qu, L.J., and Deng, X.W. (2002). Two interacting bZIP proteins are direct targets of COP1-mediated control of light-dependent gene expression in Arabidopsis. Genes Dev 16, 1247–1259.

Izawa, T., Oikawa, T., Tokutomi, S., Okuno, K., and Shimamoto, K. (2000). Phytochromes confer the photoperiodic control of flowering in rice (a short-day plant). Plant J 22, 391–399.

Keuskamp, D.H., Pollmann, S., Voesenek, L.A., Peeters, A.J., and Pierik, R. (2010). Auxin transport through PIN-FORMED 3 (PIN3) controls shade avoidance and fitness during competition. Proc Natl Acad Sci U S A 107, 22740–22744.

Klose, C. (2019). In vivo spectroscopy. Methods Mol Biol 2026, 113–120.

Kohchi, T., Mukougawa, K., Frankenberg, N., Masuda, M., Yokota, A., and Lagarias, J.C. (2001). The Arabidopsis HY2 gene encodes phytochromobilin synthase, a ferredoxin-dependent biliverdin reductase. Plant Cell 13, 425–436.

Koller, D. (1957). Germination-regulating mechanisms in some desert seeds. IV. Atriplex dimorphostegia Kar. et Kir. Ecology 38, 1–13.

Konomi, K., Abe, H., and Furuya, M. (1987). Changes in the content of phytochrome I and II apoproteins in embryonic axes of pea seeds during imbibition. Plant & Cell Physiology 28, 1443–1451.

Koornneef, M., Rolff, E., and Spruit, C.J.P. (1980). Genetic control of light-inhibited hypocotyl elongation in Arabidopsis thaliana (L.) Heynh. Zeitschrift für Pflanzenphysiologie 100, 147–160.

Koornneef, M., Cone, J.W., Dekens, R.G., O’Herne-Robers, E.G., Spruit, C.J.P., and Kendrick, R.E. (1985). Photomorphogenic responses of long hypocotyl mutants of tomato. Journal of Plant Physiology 120, 153–165.

Koren, S., Walenz, B.P., Berlin, K., Miller, J.R., Bergman, N.H., and Phillippy, A.M. (2017). Canu: scalable and accurate long-read assembly via adaptive k-mer weighting and repeat separation. Genome Res 27, 722–736.

Kraepiel, Y., Jullien, M., Cordonnier-Pratt, M.M., and Pratt, L. (1994). Identification of two loci involved in phytochrome expression in Nicotiana plumbaginifolia and lethality of the corresponding double mutant. Mol Gen Genet 242, 559–565.

Lai, L.M., Chen, L.J., Jiang, L.H., Zhou, J.H., Zheng, Y.R., and Shimizu, H. (2016). Seed germination of seven desert plants and implications for vegetation restoration. Aob Plants 8, plw031.

Lamparter, T., Esch, H., Cove, D., Hughes, J., and Hartmann, E. (1996). Aphototropic mutants of the moss *Ceratodon purpureus* with spectrally normal and with spectrally dysfunctional phytochrome. Plant, Cell & Environment 19, 560–568.

Lenser, T., Graeber, K., Cevik, Z.S., Adiguzel, N., Donmez, A.A., Grosche, C., Kettermann, M., Mayland-Quellhorst, S., Merai, Z., Mohammadin, S., Nguyen, T.P., Rumpler, F., Schulze, C., Sperber, K., Steinbrecher, T., Wiegand, N., Strnad, M., Mittelsten Scheid, O., Rensing, S.A., Schranz, M.E., Theissen, G., Mummenhoff, K., and Leubner-Metzger, G. (2016). Developmental control and plasticity of fruit and seed dimorphism in *Aethionema arabicum*. Plant Physiology 172, 1691–1707.

Lescot, M., Déhais, P., Thijs, G., Marchal, K., Moreau, Y., Van de Peer, Y., Rouzé, P., and Rombauts, S. (2002). PlantCARE, a database of plant cis-acting regulatory elements and a portal to tools for *in silico* analysis of promoter sequences. Nucleic Acids Res 30, 325–327.

Li, J., Li, G., Wang, H., and Wang Deng, X. (2011). Phytochrome signaling mechanisms. Arabidopsis Book 9, e0148.

Mahawar, L., and Shekhawat, G.S. (2018). Haem oxygenase: A functionally diverse enzyme of photosynthetic organisms and its role in phytochrome chromophore biosynthesis, cellular signalling and defence mechanisms. Plant Cell Environ 41, 483–500.

Mancinelli, A.L. (1994). The physiology of phytochrome action. In Photomorphogenesis in Plants, R.E. Kendrick and G.H.M. Kronenberg, eds (Dordrecht, The Netherlands: Kluwer Academic Publishers), pp. 211–269.

Mérai, Z., Graeber, K., Wilhelmsson, P., Ullrich, K.K., Arshad, W., Grosche, C., Tarkowská, D., Turečková, V., Strnad, M., Rensing, S.A., Leubner-Metzger, G., and Mittelsten Scheid, O. (2019). Aethionema arabicum: a novel model plant to study the light control of seed germination. J Exp Bot 70, 3313–3328.

Nagatani, A., Reed, J.W., and Chory, J. (1993). Isolation and initial characterization of Arabidopsis mutants that are deficient in phytochrome A. Plant Physiol 102, 269–277.

Neff, M.M., and Chory, J. (1998). Genetic interactions between phytochrome A, phytochrome B, and cryptochrome 1 during Arabidopsis development. Plant Physiol 118, 27–35.

Oh, E., Yamaguchi, S., Hu, J.H., Yusuke, J., Jung, B., Paik, I., Lee, H.S., Sun, T.P., Kamiya, Y., and Choi, G. (2007). PIL5, a phytochrome-interacting bHLH protein, regulates gibberellin responsiveness by binding directly to the GAI and RGA promoters in Arabidopsis seeds. Plant Cell 19, 1192–1208.

Pons, T.L. (2000). Seed responses to light. In Seeds: the ecology of regeneration in plant communities, M. Fenner, ed (CAB International), pp. 237–260.

Quail, P.H. (1997). The phytochromes: a biochemical mechanism of signaling in sight? Bioessays 19, 571–579.

Quail, P.H., Boylan, M.T., Parks, B.M., Short, T.W., Xu, Y., and Wagner, D. (1995). Phytochromes: photosensory perception and signal transduction. Science 268, 675–680.

Rancurel, C., van Tran, T., Elie, C., and Hilliou, F. (2019). SATQPCR: Website for statistical analysis of real-time quantitative PCR data. Mol Cell Probes 46, 101418.

Rizzini, L., Favory, J.J., Cloix, C., Faggionato, D., O’Hara, A., Kaiserli, E., Baumeister, R., Schäfer, E., Nagy, F., Jenkins, G.I., and Ulm, R. (2011). Perception of UV-B by the Arabidopsis UVR8 protein. Science 332, 103–106.

Saatkamp, A., Cochrane, A., Commander, L., Guja, L.K., Jimenez-Alfaro, B., Larson, J., Nicotra, A., Poschlod, P., Silveira, F.A.O., Cross, A.T., Dalziell, E.L., Dickie, J., Erickson, T.E., Fidelis, A., Fuchs, A., Golos, P.J., Hope, M., Lewandrowski, W., Merritt, D.J., Miller, B.P., Miller, R.G., Offord, C.A., Ooi, M.K.J., Satyanti, A., Sommerville, K.D., Tangney, R., Tomlinson, S., Turner, S., and Walck, J.L. (2019). A research agenda for seed-trait functional ecology. New Phytol 221, 1764–1775.

Sawers, R.J., Linley, P.J., Gutierrez-Marcos, J.F., Delli-Bovi, T., Farmer, P.R., Kohchi, T., Terry, M.J., and Brutnell, T.P. (2004). The Elm1 (ZmHy2) gene of maize encodes a phytochromobilin synthase. Plant Physiol 136, 2771–2781.

Schulz, O.P., and Klein, R.M. (1963). Effects of visible and ultraviolet radiation on the germination of *Phacelia tanacetifolia*. Am J Bot 50, 430–434.

Seo, M., Nambara, E., Choi, G., and Yamaguchi, S. (2009). Interaction of light and hormone signals in germinating seeds. Plant Molecular Biology 69, 463–472.

Seo, M., Hanada, A., Kuwahara, A., Endo, A., Okamoto, M., Yamauchi, Y., North, H., Marion-Poll, A., Sun, T.P., Koshiba, T., Kamiya, Y., Yamaguchi, S., and Nambara, E. (2006). Regulation of hormone metabolism in Arabidopsis seeds: phytochrome regulation of abscisic acid metabolism and abscisic acid regulation of gibberellin metabolism. Plant Journal 48, 354–366.

Shinomura, T., Nagatani, A., Chory, J., and Furuya, M. (1994). The induction of seed germination in *Arabidopsis thaliana* is regulated principally by phytochrome B and secondarily by phytochrome A. Plant Physiol 104, 363–371.

Shinomura, T., Nagatani, A., Hanzawa, H., Kubota, M., Watanabe, M., and Furuya, M. (1996). Action spectra for phytochrome A- and B-specific photoinduction of seed germination in *Arabidopsis thaliana*. Proc Natl Acad Sci U S A 93, 8129–8133.

Shropshire, W., Klein, W.H., and Elstad, V.B. (1961). Action spectra of photomorphogenic induction and photoinactivation of germination in *Arabidopsis thaliana*. Plant & Cell Physiology 2, 63–69.

Stawska, M., and Oracz, K. (2015). Network of signal transduction pathways mediated by phytochromes, cryptochromes and regulators of growth and development in seed biology. Postepy Biol Komorki 42, 687–706.

Stawska, M., and Oracz, K. (2019). phyB and HY5 are involved in the blue light-mediated alleviation of dormancy of Arabidopsis seeds possibly via the modulation of expression of genes related to light, GA, and ABA. Int J Mol Sci 20.

Takaki, M. (2001). New proposal of classification of seeds based on forms of phytochrome instead of photoblastism. Revista Brasileira de Fisiologia Vegetal 13, 104–108.

Tanno, N. (1983). Blue light induced inhibition of seed germination: The necessity of the fruit coats for the blue light response. Physiologia Plantarum 58, 18–20.

Terry, M.J. (1997). Phytochrome chromophore-deficient mutants. Plant, Cell & Environment 20, 740–745.

Terry, M.J., Linley, P.J., and Kohchi, T. (2002). Making light of it: the role of plant haem oxygenases in phytochrome chromophore synthesis. Biochem Soc Trans 30, 604–609.

Thanos, C.A., and Mitrakos, K. (1992). Watermelon seed germination. Osmomanipulation of photosensitivity. Seed Science Research 2, 163–168.

Thanos, C.A., Georghiou, K., and Delipetrou, P. (1994). Photoinhibition of seed germination in the maritime plant *Matthiola tricuspidat*a. Ann Bot-London 73, 639–644.

Thanos, C.A., Georghiou, K., Douma, D.J., and Marangaki, C.J. (1991). Photoinhibition of seed germination in Mediterranean maritime plants. . Ann Bot-London 68, 469–475.

Turecková, V., Novák, O., and Strnad, M. (2009). Profiling ABA metabolites in Nicotiana tabacum L. leaves by ultra-performance liquid chromatography-electrospray tandem mass spectrometry. Talanta 80, 390–399.

Urbanová, T., Tarkowská, D., Novák, O., Hedden, P., and Strnad, M. (2013). Analysis of gibberellins as free acids by ultra performance liquid chromatography-tandem mass spectrometry. Talanta 112, 85–94.

van Tuinen, A., Hanhart, C.J., Kerckhoffs, L.H.J., Nagatani, A., Boylan, M.T., Quail, P.H., Kendrick, R.E., and Koornneef, M. (1996). Analysis of phytochrome-deficient yellow-green-2 and aurea mutants of tomato. Plant Journal 9, 173–182.

Vandelook, F., Newton, R.J., and Carta, A. (2018). Photophobia in Lilioid monocots: photoinhibition of seed germination explained by seed traits, habitat adaptation and phylogenetic inertia. Ann Bot-London 121, 405–413.

Venable, D.L., and Lawlor, L. (1980). Delayed germination and dispersal in desert annuals: Escape in space and time. Oecologia 46, 272–282.

Venable, D.L., and Brown, J.S. (1988). The selective interactions of dispersal, dormancy, and seed size as adaptations for reducing risk in variable environments. The American Naturalist 131, 360–384.

Vleeshouwers, I.M., Bouwmeester, H.J., and Karssen, C.M. (1995). Redefining seed dormancy: An attempt to integrate physiology and ecology. Journal of Ecology 83, 1031–1037.

Weller, J.L., Murfet, I.C., and Reid, J.B. (1997). Pea mutants with reduced sensitivity to far-red light define an important role for phytochrome A in day-length detection. Plant Physiol 114, 1225–1236.

Weller, J.L., Terry, M.J., Rameau, C., Reid, J.B., and Kendrick, R.E. (1996). The phytochrome-deficient pcd1 mutant of pea is unable to convert heme to biliverdin IX alpha. Plant Cell 8, 55–67.

Wu, T.D., and Watanabe, C.K. (2005). GMAP: a genomic mapping and alignment program for mRNA and EST sequences. Bioinformatics 21, 1859–1875.

Yang, L., Liu, S., and Lin, R. (2020). The role of light in regulating seed dormancy and germination. J Integr Plant Biol 62, 1310–1326.

Yaniv, Z., and Mancinelli, A.L. (1968). Phytochrome and seed germination. IV. Action of light sources with different spectral energy distribution on the germination of tmato seeds. Plant Physiol 43, 117–120.

